# Direct and High-Throughput Assays for Human Cell Killing through Trogocytosis by *Entamoeba histolytica*

**DOI:** 10.1101/768069

**Authors:** Akhila Bettadapur, Katherine S. Ralston

## Abstract

*Entamoeba histolytica* is a microbial eukaryote and causative agent of the diarrheal disease amoebiasis. Pathogenesis is associated with profound damage to human tissues, and treatment options are limited. We discovered that amoebae attack and kill human cells through a cell-nibbling process that we named trogocytosis (*trogo-:* nibble). Trogocytosis is likely to underlie tissue damage during infection and it represents a potential target for therapeutic intervention, although the mechanism is still unknown. Assays in current use to analyze trogocytosis by amoebae have not been amenable to studying different types of human cells, or to high-throughput analysis. Here, we developed two complementary assays to measure trogocytosis by quantifying human cell viability, both of which can be used for suspension and adherent cells. The first assay uses CellTiterGlo, a luminescent readout for cellular ATP levels, as a proxy for cell viability. We found that the CellTiterGlo signal is proportional to the quantity of viable cells, and can be used to detect death of human cells after co-incubation with amoebae. We established a second assay that is microscopy-based and uses two fluorescent stains to directly differentiate live and dead human cells. Both assays are simple and inexpensive, can be used with suspension and adherent human cell types, and are amenable to high-throughput approaches. These new assays are tools to improve understanding of amoebiasis pathogenesis.

## 1. Introduction

*Entamoeba histolytica* is the causative agent of amoebiasis. During infection, the actively replicating trophozoite (amoeba) form colonizes the large intestine. Symptoms range from asymptomatic infection, diarrhea, bloody diarrhea, to fatal extraintestinal abscesses. The species name (*histo*-: tissue; *lytic*-: dissolve) refers to the capacity of the amoeba to damage human tissues. However, it is still unclear how amoebae invade and damage tissues. Virulence factors include the amoeba surface galactose and N-acetylgalactosamine (Gal/GalNAc)-inhibitable lectin that mediates attachment to human cells and other substrates (Petri, 2002) and cysteine proteases that degrade a variety of human substrates (*e.g.,* Reed, 1995, Lidell, 2006, Thibeaux, 2012). In addition to these factors, the contact-dependent human-cell killing activity of *E. histolytica* (Ravdin, 1980a, Ravdin, 1981) is likely to be a major contributor to human tissue damage.

While it has been under investigation for many years, the mechanism by which amoebae kill human cells was previously unclear (Ralston, 2011). We defined that amoebae kill human cells *via* trogocytosis (*trogo-:* nibble) (Ralston, 2014). Amoebae attach to human cells and then physically extract “bites” of human cell membrane, cytoplasm and organelles, which eventually leads to human cell death. Amoebic trogocytosis requires engagement of the Gal/GalNAc lectin, actin rearrangements, PI3K and EhC2PK signaling (Ralston, 2014). Amoebic trogocytosis is necessary for invasion of *ex vivo* intestinal tissue, underlining its relevance to pathogenesis (Ralston, 2014). Trogocytosis might be evolutionarily-conserved (Ralston, 2015), therefore, studying this process in *E. histolytica* may give insight into eukaryotic trogocytosis, in addition to a better understanding of the pathogenesis of amoebiasis.

In order to better understand trogocytosis and its contribution to disease, there is a need for cell death assays that are accurate, practical and that can be applied to a variety of human cell types. Assaying human cell killing by *E. histolytica* is inherently challenging since readouts must specifically measure the viability of the human cells when they are mixed together with amoebae. For the greatest utility, assays must directly measure human cell viability, and readouts must be quantitative. While amoebae can kill essentially any human cell type (Ravdin, 1980b), most studies have focused on either monolayers or suspension cultures, but not both, since they are typically not amenable to the same assays. Thus, there is a need for flexible assays that can be applied to both monolayers and suspension cells. Although previously used assays have been important in advancing understanding of cell killing by amoebae, it is important to recognize their limitations and to develop new assays as newer technologies become available.

Assays that have been used can be broken down into membrane permeabilization, monolayer disruption, and apoptosis assays. Membrane permeabilization assays detect intracellular components that are released into the culture supernatant by dead cells. In these assays, amoebae are co-incubated with human cells, and the supernatant is measured. There are some technical and practical limitations to the lactate dehydrogenase (LDH) release and Chromium-51 (^51^Cr) release assays that have been used. LDH assays (*e.g.,* Li, 1994, Marie, 2012) generally do not directly measure LDH and instead use the NAD cofactor to catalyze a reporter reaction (Riss, 2019). This means that other enzymatic activities in the culture supernatant that also use NAD as a cofactor can be problematic. By contrast, the ^51^Cr release assay specifically measures host cell lysis, since in this assay, host cells are pre-labeled with ^51^Cr, and after incubation with amoebae, ^51^Cr in the culture supernatant is measured (*e.g.,* Saffer, 1991, Huston, 2001). However, a practical limitation is that this assay requires the use of a radioisotope.

Monolayer disruption assays have been used in many studies, but the major limitation is that monolayer disruption cannot be directly attributed to cell killing since amoebic cysteine protease activity disrupts monolayers (Tillack, 2006). In monolayer disruption assays, amoebae are incubated with host cell monolayers, and after washing, the remaining cells are stained with methylene blue (*e.g.,* Bracha, 1984, Teixeira, 2012). The amount of methylene blue is compared to control monolayers that were incubated without amoebae, to infer how many cells have been released. Trypan blue staining has also been used to stain dead cells remaining in the monolayer (*e.g.,* Ravdin, 1985, Bracha, 1999). However, since amoebic proteases cause disruption of monolayers (Tillack, 2006), neither version of this assay directly measures cell killing.

Finally, apoptosis assays have been used to study cell killing by amoebae (Seydel, 1998, Huston, 2001). In these assays, care must be taken to include controls that ensure the readout is specific to apoptosis. For example, DNA laddering can occur in other modes of cell death besides canonical apoptosis, and thus is not indicative of apoptotic cell death (Kroemer, 2008). As another example, annexin V staining to detect exposed phosphatidylserine must be combined with cell permeability stains like propidium iodide, in order to ensure that phosphatidylserine exposure is not simply the result of membrane damage. Notably, phosphatidylserine exposure is not a universal feature of apoptosis (Galluzzi, 2018). It is also important to note that apoptosis assays capture markers of apoptosis in dying cells, which differs from other cell death assays that measure cell death after it has occurred. Notably, in some cases, apoptosis can be reversed (Tang, 2012). Thus, cell death is inferred by these assays, but not directly measured.

To enable quantitative cell death measurements, we previously developed an assay using imaging flow cytometry (Ralston, 2014). In this assay, amoebae and human cells are fluorescently labeled, allowing for trogocytosis to be directly measured, and Live/Dead Violet is used to stain dead (permeable) cells (Ralston, 2014, Gilmartin, 2017, Miller, 2019). This assay allows for automated analysis of thousands of images per sample, but is limited in practicality since imaging flow cytometers are not widely available. This assay is more easily applied to study suspension cells, since cells must be in suspension during image acquisition; thus, imaging flow cytometry is limited in flexibility.

To address the limitations in practicality and flexibility of existing cell death assays, here we developed two complementary high-throughput assays for human cell death. We show that CellTiterGlo, a luminescent readout for cellular ATP levels, can be used as a proxy for human cell viability. We also develop a confocal microscopy-based assay with fluorescent stains to quantitatively differentiate live and dead human cells. Both assays are simple and inexpensive, and they can be used with suspension and adherent human cell types.

## 2. Materials and Methods

### 2.1 Cell culture

*E. histolytica* HM1:IMSS (ATCC) trophozoites were cultured in TYI-S-33 media, supplemented with 15% heat inactivated Adult Bovine Serum (Gemini Bio Products), 80 units/mL penicillin and streptomycin (Gibco), and 2.3% Diamond Vitamin solution 80 Tween 40x (Sigma Aldrich), at 35°C. Amoebae were harvested when flasks were approximately 80% confluent. Human Jurkat T cells, clone E6-1 (ATCC), were cultured in RPMI Medium 1640 with L-Glutamine and without Phenol Red (Gibco), supplemented with 10 mM Hepes (Affymetrix), 100 units/mL penicillin and streptomycin (Gibco) and 10% heat inactivated Fetal Bovine Serum (Gibco), at 37°C with 5% CO_2_. Cells were harvested at approximately 1×10^6^ cells/mL. Human Caco-2 colon epithelial cells, HTB-37 (ATCC), were cultured in MEM Medium (ATCC), supplemented with 20% Fetal Bovine Serum (Gibco), at 37°C with 5% CO_2_. Cells were passaged using 0.25% (w/v) Trypsin – 0.53 mM EDTA solution when 80-100% confluent.

### 2.2 Knockdown mutants

The EhROM1 silencing construct was generated by Morf, *et al.* (Morf, 2013), and contains 132 base pairs of the trigger gene (EHI_048600) fused to the first 537 base pairs of EhROM1 (EHI_197460). This plasmid, or a corresponding vector control, was transfected into amoebae using Attractene transfection reagent (Qiagen), and then stable transfectants were selected and maintained with Geneticin at 6 μg/mL (Invitrogen). Clonal lines were obtained by limiting dilution, and silencing was confirmed using RT-PCR (Miller, 2019). A single clonal line was used for experiments.

### 2.3 CellTiterGlo Assay

For experiments using Jurkat cells, amoebae and Jurkat cells were first washed in fresh TYI media. For the initial titration experiments (Fig. 1A – 1B), amoebae and Jurkat cells were resuspended to 2×10^6^ and 1×10^7^ cells/mL, respectively. For Cytochalasin D experiments, amoebae were first washed in fresh TYI media and pretreated with 20 nM Cytochalasin D from *Zygosporium mansonii* (Sigma Aldrich) or an equivalent volume of DMSO for 1 hour at 35°C. Cytochalasin D, or DMSO, was maintained at the same concentration when amoebae were subsequently co-incubated with Jurkat cells. For sugar inhibition experiments, amoebae were resuspended in fresh TYI media with no supplementation, 100 mM galactose (Sigma Aldrich), or 100 mM mannose (Sigma Aldrich). For the initial co-incubation assays (Fig. 1C – 1D and Fig. S1), amoebae and Jurkat cells were resuspended to 4×10^5^ and 2×10^6^ cells/mL, respectively, to create a co-incubation ratio of 1 amoeba: 5 Jurkat cells. For all other co-incubation assays, amoebae and Jurkat cells were resuspended to 4×10^5^ and 8×10^6^ cells/mL respectively, to create a co-incubation ratio of 1:20.

**Figure 1:**
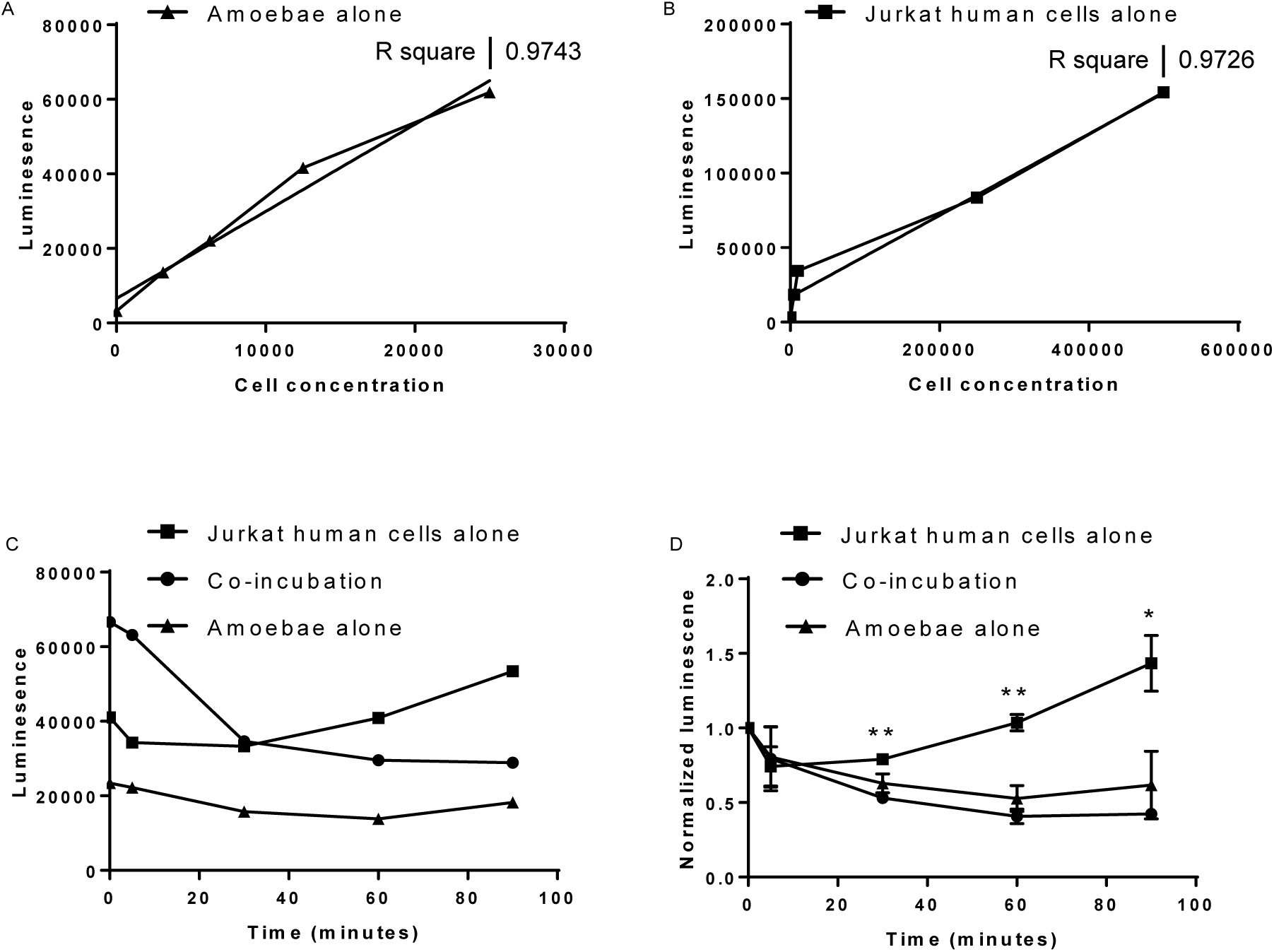
CellTiterGlo can be used to assay Jurkat cell killing by amoebae. **(A)** A dilution series of amoebae, or **(B)** human Jurkat T cells was assayed using CellTiterGlo. Best fit lines and R^2^ values are shown. CellTiterGlo signal correlates with the number of cells per well. Data represent the average values of two replicate wells for each cell concentration, and are representative of 3 independent experiments. **(C)** Amoebae were co-incubated (filled circles) with Jurkat cells at a 1:5 ratio, or amoebae (filled triangles) and Jurkat cells (filled squares) were incubated separately as controls. Data represent the average values of two replicate wells for each sample, from one experiment. **(D)** Data from 2 independent experiments performed as in Panel C were normalized to the value of each sample at Time = 0. There were statistically significant differences between the co-incubation and Jurkat alone samples, as indicated.

50 μL of amoebae or Jurkat cells were plated in 96 well plates (Corning 3603) either individually, with 50 μL of TYI media, or together. Plates were placed in an anerobic GasPak (BD) and incubated at 35°C for the appropriate time. At each time point, a plate was removed from the incubator and left at 25°C for 10 minutes to equilibrate. 100 μL of CellTiterGlo solution was added to each well, using a multichannel pipette. Plates were then incubated at 25°C for 10 minutes, with rocking, and then luminescence was detected using a 1 second exposure on a plate reader (PerkinElmer 2030 Victor). Two wells for each condition were averaged to generate one value per condition, and at least three experiments were performed independently on different days. Both raw data from individual experiments and normalized data from multiple independent experiments are presented in the figures. For normalization, data from multiple independent experiments were normalized to the T = 0 time point for each sample.

For experiments using Caco-2 cells, 18-24 hours prior to performing the experiment, 100 μL of Caco-2 cells were plated in to three 96-well plates (Corning 3603) at a concentration of 2.6×10^5^ cells/mL. On the day of the CellTiterGlo assay, wells containing Caco-2 cells were gently washed with 200 μL of fresh TYI, twice. Amoebae were washed in fresh TYI media and resuspended to 2.6×10^4^ cells/mL. 100 μL of amoebae were added to wells containing Caco-2 cells to create an approximate co-incubation ratio of 1 amoeba: 10 Caco-2 cells. Plates were incubated and treated with CellTiterGlo as described above. Three wells of amoebae alone, six wells of Caco-2 cells alone, and six wells of co-incubated samples were averaged to obtain one value per condition, and three experiments were performed independently on different days.

### 2.4 Dual-Stain Microscopy Assay

For experiments using Jurkat cells, amoebae were washed in fresh TYI media and pre-treated with 20 nM Cytochalasin D from *Zygosporium mansonii* (Sigma Aldrich) or an equivalent volume of DMSO for 1 hour at 35°C. Cytochalasin D, or DMSO, was maintained at the same concentration when amoebae were subsequently co-incubated with Jurkat cells. Jurkat cells were pre-labeled with Hoechst 33342 (Invitrogen) at 5 μg/ml for 30 minutes at 37°C. Amoebae and Jurkat cells were then washed in M199S (Gibco M199 with Earle’s Salts, L-Glutamine, 2.2 g/L Sodium Bicarbonate and without Phenol Red, and supplemented with 5.7 mM L-cysteine (Sigma-Aldrich), 25 mM HEPES (Sigma-Aldrich) and 0.5% bovine serum albumin (Gemini Bio-Products)). Amoebae and Jurkat cells were then resuspended to 2×10^5^ and 1×10^6^ cells/mL, respectively in M199S containing 20 nM SYTOX green (Thermo), and 20 nM Cytochalasin D or an equivalent volume of DMSO. 1 mL of each cell type was added to 35 mm glass bottom petri dishes containing a N° 1.5 coverglass (MatTek). Petri dishes were warmed to 35°C for 15 minutes before use. Cells were co-incubated at 35°C for 60 minutes before confocal microscopy imaging. Cells were imaged using a stage warmer set to 35°C on either an Intelligent Imaging Innovations hybrid spinning disk confocal microscope or an Olympus FV1000 laser point-scanning confocal microscope. Two experiments were performed independently on different days, and 350-500 human cells were counted for each condition.

For experiments using Caco-2 cells, Caco-2 cells were cultured on collagen-coated (5 μg/cm^2^ Collagen I Rat Tail, Gibco) glass bottom petri dishes containing a N° 1.5 coverglass (MatTek). Experiments were performed when cells were ∼80% confluent. Caco-2 cells were pre-labeled by incubation in 2 mL of M199S containing Hoechst 33342 at 5 μg/ml for 30 minutes at 37°C. Amoebae and Caco-2 cells were then washed in M199S. Amoebae were resuspended to 1×10^5^ cells/mL in M199S containing 20 nM SYTOX green, and 2 mL of amoebae were then added to each plate containing Caco-2 cells. Plates were then incubated at 35°C. Cells were imaged using a stage warmer set to 35°C on an Olympus FV1000 laser point-scanning confocal microscope.

### 2.5 Statistical Analysis

GraphPad Prism was used to calculate best fit line and R^2^ values, and for student’s unpaired *t* test statistical analysis. Mean values and standard deviations are shown in the figures, with *t* test values reported as follows: ns = P > 0.05, * = P < 0.05, ** = P < 0.01, *** = P < 0.001, **** = P < 0.0001.

## 3. Results

CellTiterGlo is a very simple, high-throughput assay for cell viability that is based on cellular ATP levels. We reasoned that since human cells are present in excess of amoebae, they should contribute to the majority of the CellTiterGlo luminescence signal in a co-incubation. We first asked whether luminescence values correlated with the number of amoebae (Fig. 1A) or human Jurkat T cells per well (Fig. 1B), and found that luminescence was correlated with cell number. Next, amoebae and Jurkat cells were co-incubated, or as controls, Jurkat human cells or amoebae were incubated alone. In these controls, the equivalent number of cells were loaded per well to correspond to the number of cells present in the co-incubation experimental condition. As anticipated, in the controls, the luminescence values were higher for human cells than for amoebae (Fig. 1C – 1D). When human cells and amoebae were co-incubated, the luminescence value initially corresponded to roughly the sum of the human cell and amoeba individual values, and then decreased over time. The luminescence values of the co-incubation were significantly lower than the values for human cells incubated alone (Fig. 1D). Reduced variability was observed when samples were incubated in an anaerobic GasPak (Fig. 1C – 1D), compared to an aerobic environment (Fig. S1), consistent with the microaerophilic metabolism of *E. histolytica.* Therefore, we concluded that human cell killing by amoebae can be quantitatively measured using CellTiterGlo.

We next asked whether this assay was sensitive to conditions that inhibit human cell killing by amoebae. Trogocytosis by *E. histolytica* requires actin rearrangements and is inhibited by treatment with cytochalasin D (Ralston, 2014). Therefore, cytochalasin D-treated or DMSO control-treated amoebae were co-incubated with Jurkat cells, and cell viability was measured using CellTiterGlo (Fig. 2A – 2B). Cytochalasin D-treated amoebae were significantly less able to kill human cells, as seen by the increased CellTiterGlo signal compared to control amoebae (Fig. 2A – 2B). For human cells or amoebae incubated alone, cytochalasin D treatment did not affect viability (Fig. S2A). We next tested if CellTiterGlo was sensitive to inhibition of the amoeba surface Gal/GalNAc lectin. Amoeba must attach to human cells in order to kill them, and this attachment is mediated by the amoebic GalNAc lectin (Petri, 2002). Galactose-treated amoebae were significantly less able to kill human cells, compared to control mannose-treated amoebae (Fig. 2C, S2B). Finally, we tested knockdown mutant amoebae deficient in a rhomboid protease, EhRom1, which has a characterized role in attachment to human cells (Baxt, 2008, Baxt, 2010). There was no significant difference in the cell killing ability between EhRom1 knockdown mutants and vector control amoebae (Fig. 2D, S2C). This is consistent with the lack of a trogocytosis defect in EhRom1 mutants (Miller, 2019). Taken together, we concluded that the CellTiterGlo assay is sensitive to the inhibition of human cell killing by amoebae.

**Figure 2:**
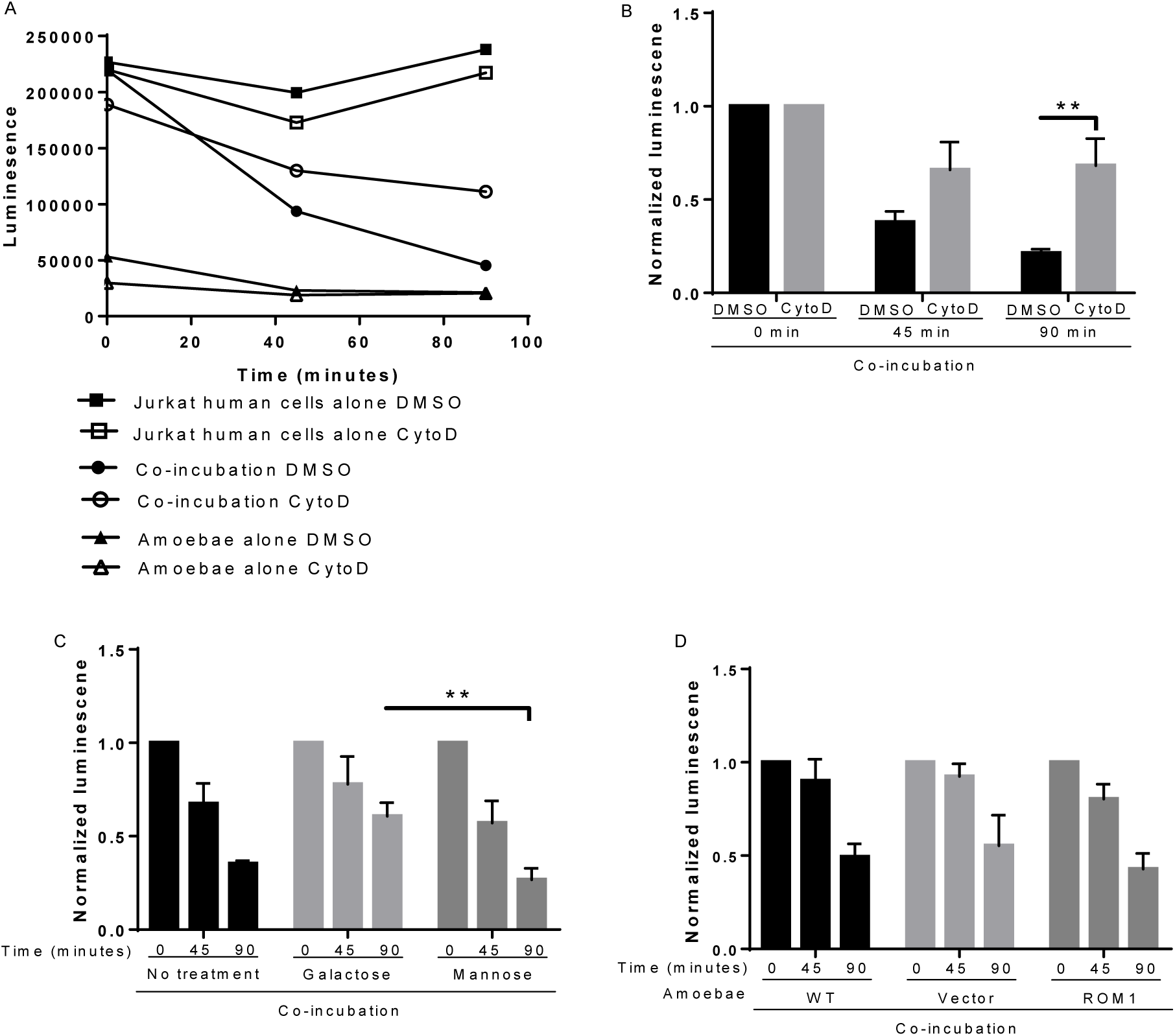
CellTiterGlo can be used to assay trogocytosis inhibition and attachment inhibition. **(A)** Amoebae and human Jurkat T cells were treated with Cytochalasin D (open symbols) or DMSO (filled symbols). Amoebae were co-incubated (circles) with Jurkat cells at a 1:20 ratio, or amoebae (triangles) and Jurkat cells (squares) were incubated separately as controls. Viability was assayed by using CellTiterGlo. Data represent the average values of two replicate wells for each sample, from one experiment. **(B)** Data from 4 independent experiments performed as in panel A were normalized to the value of each sample at Time = 0. **(C)** Amoebae and Jurkat cells were incubated in media containing galactose, mannose, or no added sugar. Cells were co-incubated at a 1:20 ratio, or incubated separately as controls. Viability was assayed by using CellTiterGlo. Data from 3 independent experiments were normalized to the value of each sample at Time = 0. **(D)** Amoebae were transfected with an EhRom1 knockdown plasmid, or a vector control plasmid. Transfectants, or wild-type non-transfected amoebae, were co-incubated with Jurkat cells at a 1:20 ratio, or incubated separately as controls. Data from 3 independent experiments were normalized to the value of each sample at Time = 0.

We next sought to extend this assay to other human cell types, since many assays for cell killing are difficult to adapt to both suspension and monolayer cells. Therefore, we adapted the CellTiterGlo assay to human Caco-2 intestinal epithelial cell monolayers. CellTiterGlo luminescence values correlated closely with the number of amoebae or Caco-2 cells per well (Fig. 3A – 3B). When Caco-2 cells and amoebae were co-incubated, the luminescence value initially corresponded to roughly the sum of the human cell and amoeba individual values (Fig. 3C). The luminescence values of the co-incubation were significantly lower than the values for human cells incubated alone (Fig. 3D). These results show that CellTiterGlo can be applied to assay killing of both suspension and monolayer cells.

**Figure 3:**
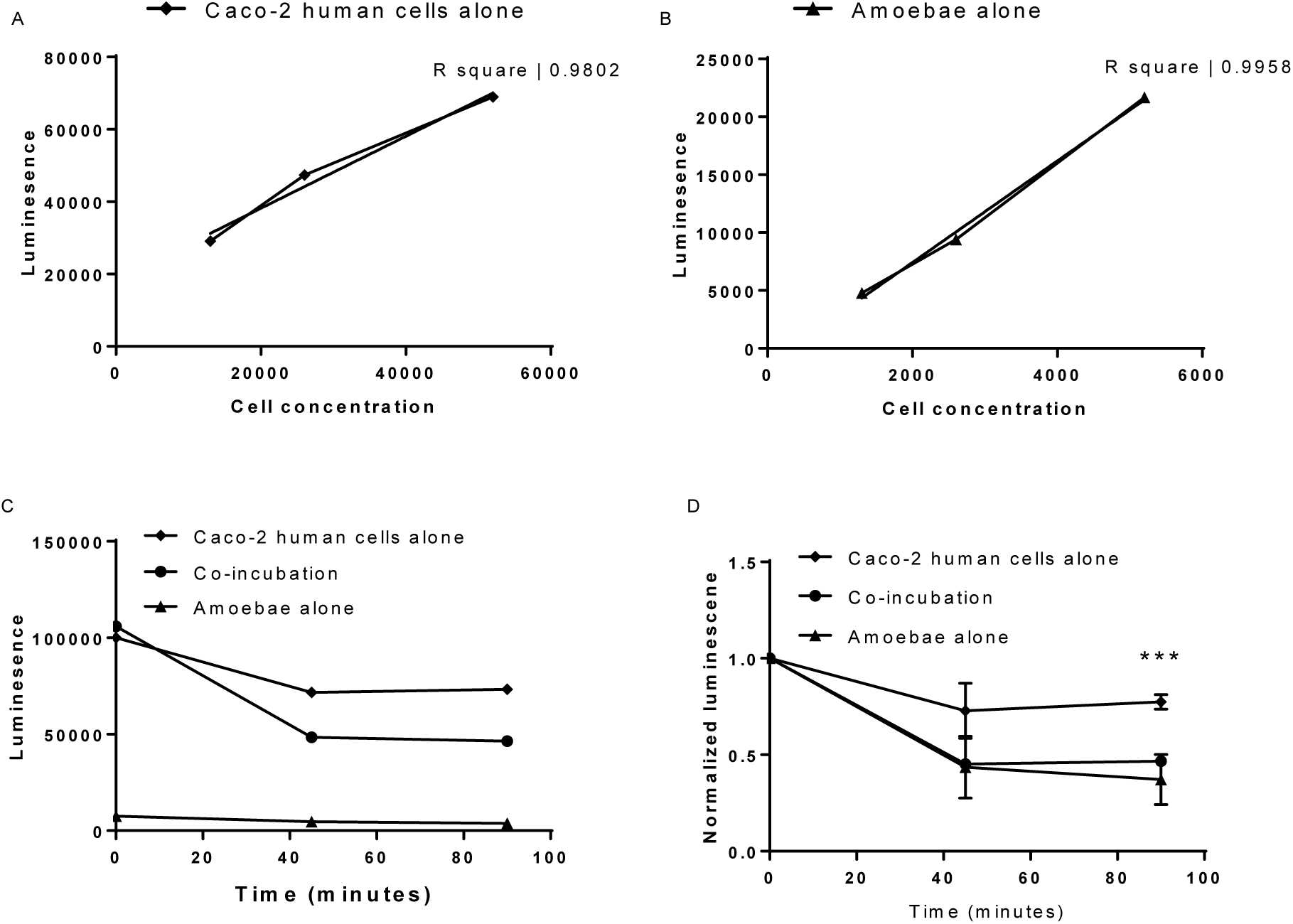
CellTiterGlo can be used to assay Caco-2 cell killing by amoebae. **(A)** A dilution series of amoebae, or **(B)** human Caco-2 intestinal epithelial cells was assayed using CellTiterGlo. Best fit lines and R^2^ values are shown. CellTiterGlo signal correlates with the number of cells per well. Data represent the average values of two replicate wells for each cell concentration, and are representative of 3 independent experiments. **(C)** Amoebae were co-incubated (filled circles) with Caco-2 cells, or amoebae (filled triangles) and Caco-2 cells (filled squares) were incubated separately as controls. Data represent the average values of two replicate wells for each sample, from one experiment. **(D)** Data from 3 independent experiments performed as in panel C were normalized to the value of each sample at Time = 0. There were statistically significant differences between the co-incubation and Caco-2 alone samples, as indicated.

We next developed a microscopy-based assay to directly measure human cell killing by amoebae. Since human cell nuclei are not internalized during amoebic trogocytosis (Ralston, 2014), we devised a strategy with two different nuclear stains to distinguish living and dead human cells. Human cell nuclei were pre-labeled with Hoechst. During co-incubation, SYTOX green was present in the media. SYTOX green is a nucleic acid stain that is excluded by living cells, but is taken up by dead cells because they have permeable membranes. Thus, live human cells are labeled only by Hoechst, while dead human cells are dual-labeled by both Hoechst and SYTOX green (Supplemental Video 1). To test this dual-stain assay, cytochalasin D treatment was used to inhibit amoebic trogocytosis. Amoebae were treated with cytochalasin D or DMSO, and co-incubated with human Jurkat T cells (Fig. 4A). Cytochalasin D-treated amoebae killed less than 2% of Jurkat cells in 60 minutes (Fig. 4B). By comparison, control amoebae killed 40% of Jurkat cells in 60 minutes. This assay provides a quantitative readout for cell killing, and it is robust enough to be amenable to imaging large fields of cells at low magnification (Fig. 4C). Finally, this dual-stain assay can be applied to Caco-2 epithelial cells (Fig. 5 and Supplemental Videos 2 – 3), demonstrating that it is versatile with respect to human cell types.

**Figure 4:**
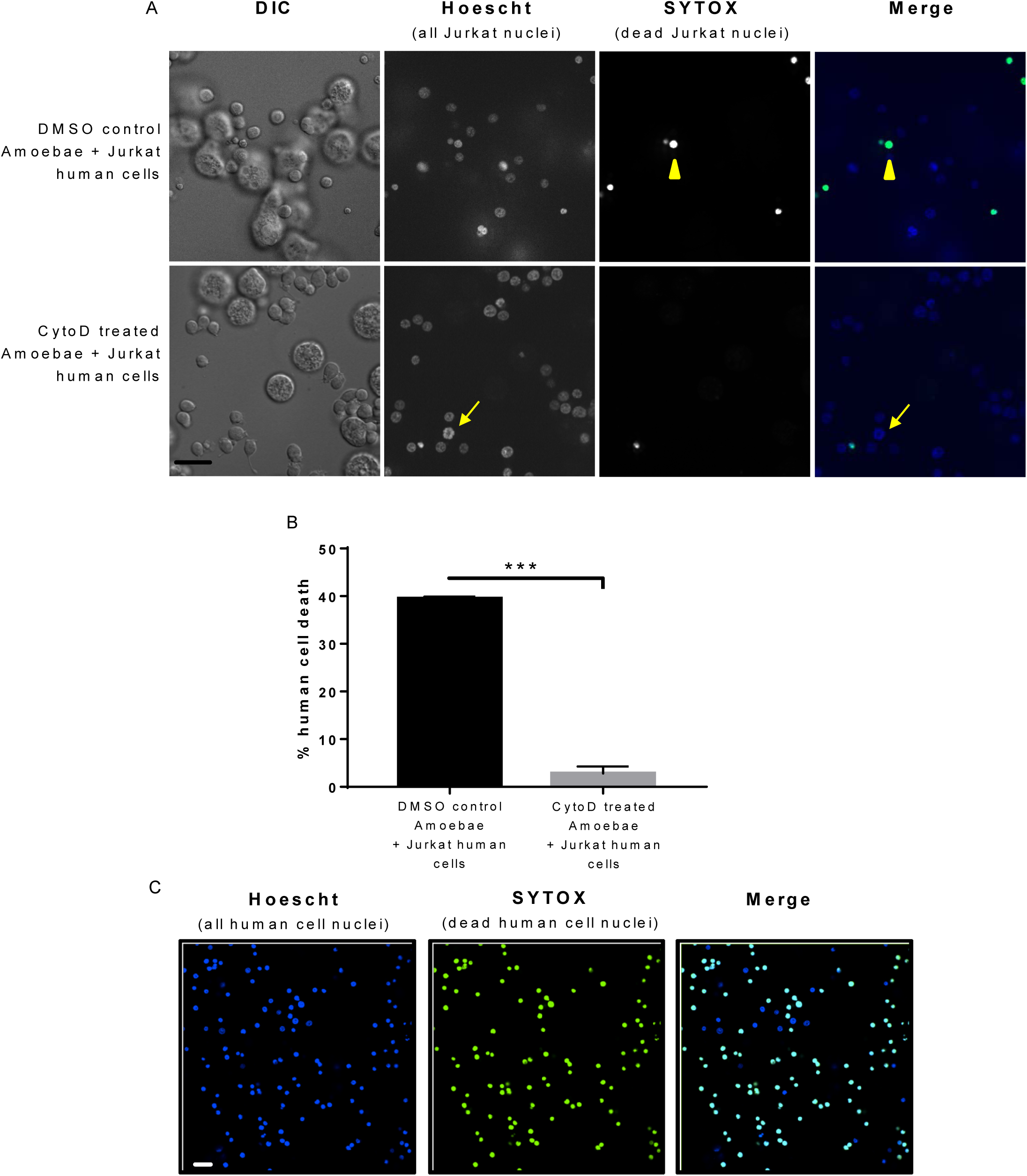
A dual-stain microscopy assay can be used to quantitatively and directly detect Jurkat cell killing by amoebae. **(A)** Amoebae and Hoechst-labeled human Jurkat T cells were treated with Cytochalasin D or DMSO, and co-incubated for 60 minutes in the presence of SYTOX green. Representative images are shown. Living human cells are labeled by Hoechst (blue), while dead human cells are labeled by both Hoechst and SYTOX green (green) and appear as turquoise in the merged image. An example of a living cell (arrow) and a dead cell (arrowhead) are indicated. Scale bar, 50 μm. **(B)** Human cell death was assayed by quantifying the number of single-stained (Hoechst) and dual-stained (Hoescht and SYTOX green) human cell nuclei, which correspond to living and dead human cells, respectively. Data are representative of 2 independent experiments. **(C)** Representative images demonstrating that the dual-stain assay can be applied to low magnification objectives, allowing for a greater number of cells to be imaged per field. Scale bar, 50 μm.

**Figure 5:**
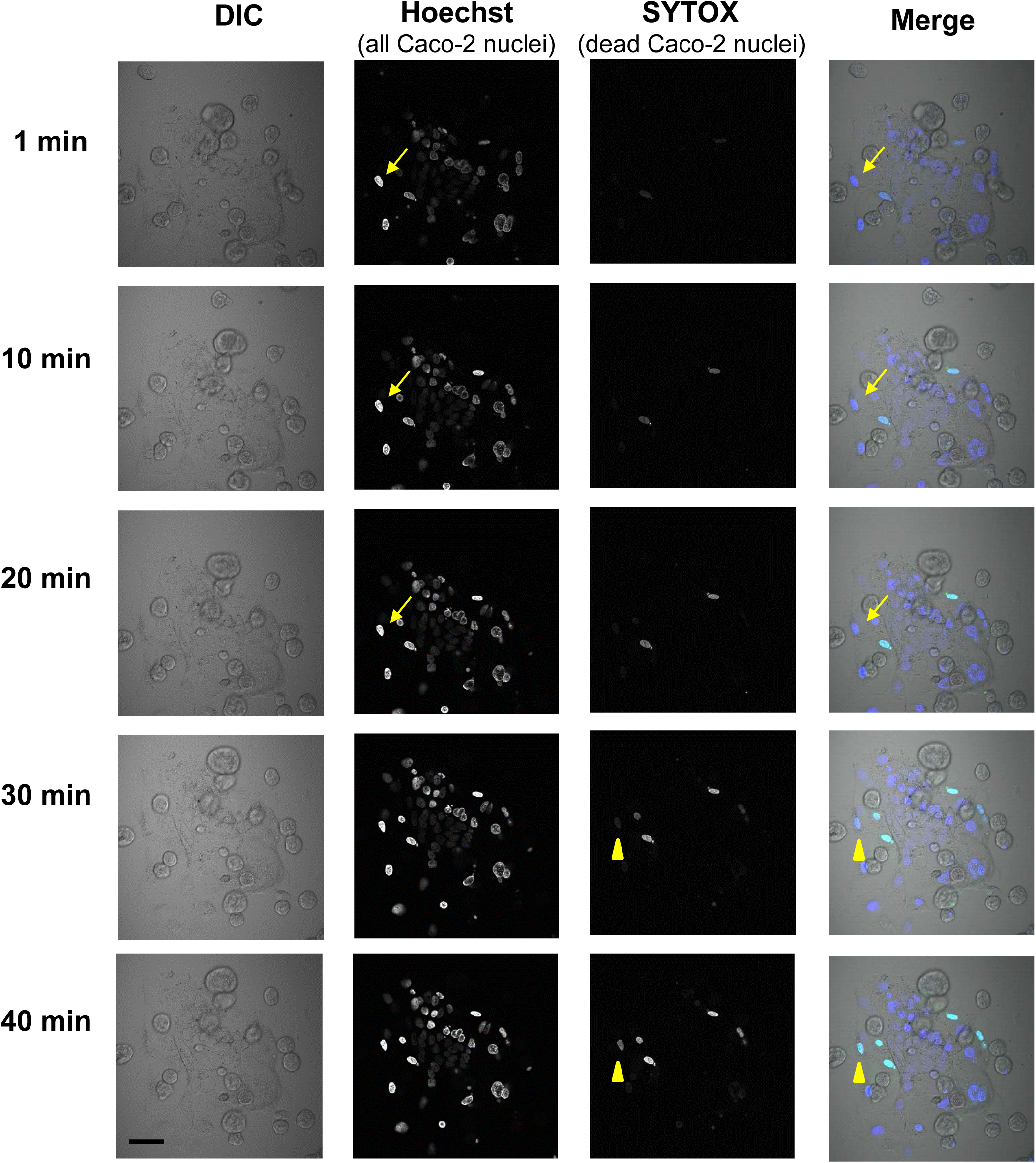
A dual-stain microscopy assay can be used to directly detect Caco-2 cell killing by amoebae. Amoebae and Hoechst-labeled human Caco-2 intestinal epithelial cells were co-incubated in the presence of SYTOX green. Living human cells are labeled by Hoechst (blue), while dead human cells are labeled by both Hoechst and SYTOX green (green) and appear as turquoise in the merged image. The arrow indicates an example of a Caco-2 cell that is initially living and labeled only by Hoechst, but is eventually killed by an amoeba, at which time it becomes labeled by SYTOX green (arrowhead). Data are representative of 2 independent experiments. Scale bar, 50 μm.

## 4. Discussion

In this study, we developed two assays for human cell killing by *E. histolytica.* The CellTiterGlo assay biochemically measures cellular ATP levels, and the dual-stain microscopy assay allows for direct visualization of human cell death with fluorescent stains. These assays complement the currently available cell death assays and bring their own unique strengths and weaknesses.

The CellTiterGlo assay is simple and practical. Only a plate reader is required for the readout. The assay requires few manipulations and no washing steps; CellTiterGlo solution is added directly to cells, and after a brief incubation, luminescence is measured on a plate reader. Since this assay is robust and requires very few steps, the procedure is amenable to high throughput screening. Indeed, CellTiterGlo has previously been used in a high throughput screen for drugs that kill *E. histolytica* (Debnath, 2012). The limitation of this assay is that it does not directly measure human cell death. Because human cells greatly outnumber amoebae in this assay, they contribute the majority of the ATP to the readout, and thus, a decrease in luminescence can be inferred to represent human cell death. Also, similar to the limitations of apoptosis assays, ATP levels are correlated with dying cells, but do not clearly define the “point of no return” when a cell is by definition, dead (Leist, 1997, Bonora, 2012). The depletion of ATP below a threshold, combined with redox alterations, has been proposed to mark the “point of no return” (Galluzzi, 2015), however, it would be difficult to infer from an assay like CellTiterGlo that the level of ATP has definitively crossed a threshold. However, the major strengths of this assay are the simplicity and adaptability to high throughput approaches. This is an area where none of the existing cell death assays are useful. Thus, we propose that this assay is most useful for initial screening of mutants or candidate inhibitors. Finally, since monolayers can be grown directly in the plates used for this assay, it is easily adaptable to both monolayers and suspension cells.

The dual-stain microscopy-based cell death assay is also simple and practical. We used confocal microscopy for imaging, but this is not necessary, as widefield fluorescence microscopes can also be used. The major strength of this assay is that it directly measures human cell death. Dead cells are labeled, thus human cell death can be directly quantified within a mixture of human cells and amoebae. Moreover, the readout for cell death in this assay is loss of membrane integrity, which is a direct marker of cells that are dead (Kroemer, 2008). Thus, this assay, together with the imaging flow cytometry assay that we developed (Ralston, 2014), represents the most direct assays available for human cell killing for *E. histolytica.* The dual-stain assay is more practical and easy to apply, since imaging flow cytometers are not widely available. Like imaging flow cytometry, the dual-stain assay can be applied to medium throughput approaches, as it could be performed by using cells in plates and by imaging on high content screening microscopes. The limitation of this assay is that the readout is not inherently quantitative, and requires counting of labeled cell nuclei. We did not develop automated image analysis, but this would be possible, and would ensure that the readout is unbiased and efficient. Because both stains label the same cellular feature, the nucleus, automated image analysis should be particularly robust. Finally, like the CellTiterGlo assay, the dual-stain microscopy assay is amenable to both monolayer and suspension cell cultures.

Together, these assays expand the repertoire of available tools for studying human cell killing by *E. histolytica.* They are particularly simple and practical, and thus we believe they are suitable for wide application. These assays also pave the way for high-throughput studies. Since cell killing by *E. histolytica* is likely to underlie disease pathogenesis, these tools are expected to allow for an improved understanding of the mechanism of disease, and may be applicable to the development of new therapeutics.

## Acknowledgments

We thank the MCB Microscopy Imaging Facility at UC Davis for use of core microscopes and technical assistance. We thank the Ralston lab for stimulating discussions and support. This work was supported by NIH grant 1R01AI146914 and a Pew Scholarship awarded to K.S.R., and the UC Davis Biotechnology Training Program fellowship awarded to A.B.

Declaration of interest: none.

---

**Supplemental Figure 1:**
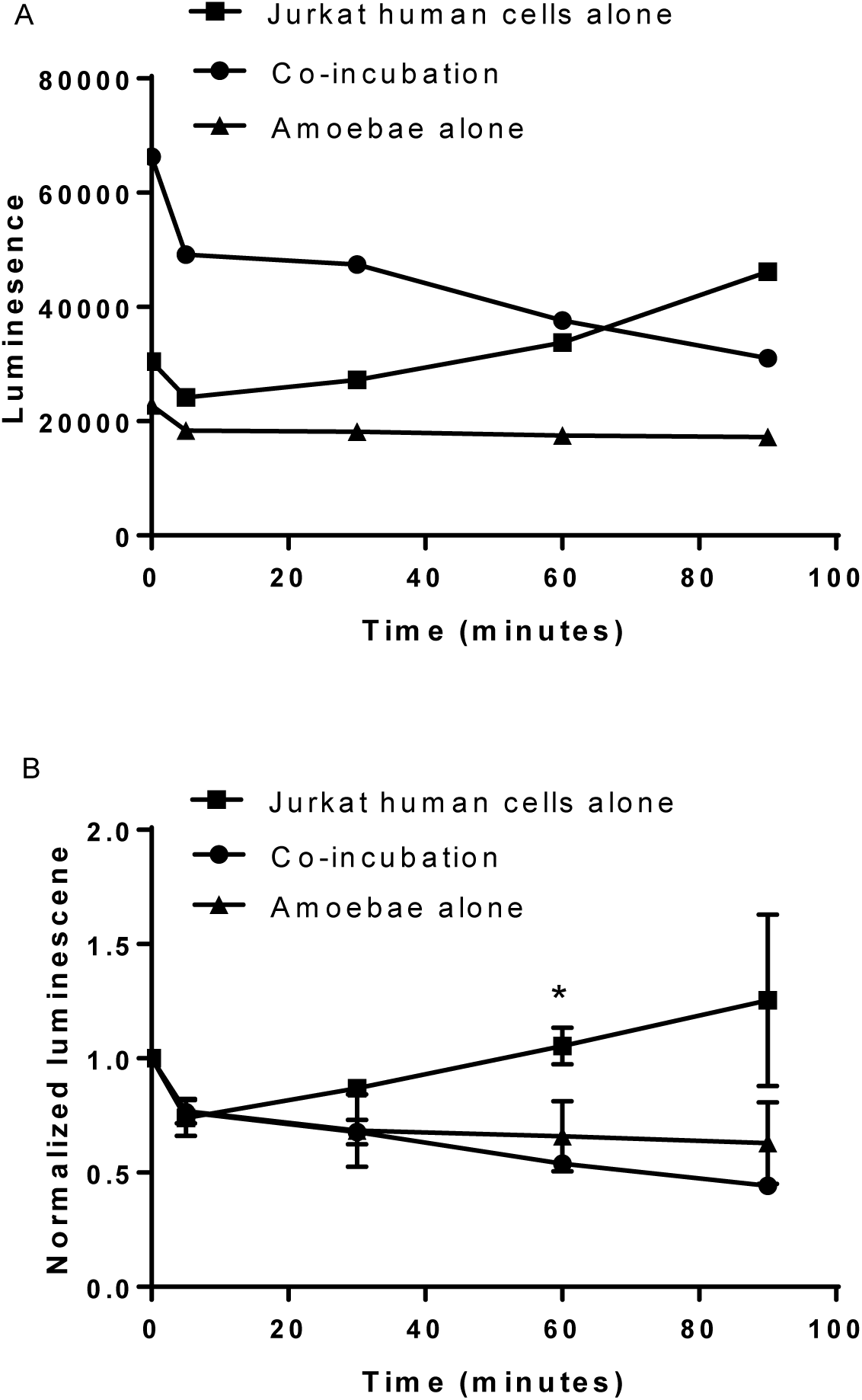
Greater variability is observed in the CellTiterGlo assay when cells are incubated without an anaerobic GasPak. **(A)** Amoebae were co-incubated (filled circles) with Jurkat cells at a 1:5 ratio, or amoebae (filled triangles) and Jurkat cells (filled squares) were incubated separately as controls. Data represent the average values of two replicate wells for each sample, from one experiment. **(B)** Data from 2 independent experiments performed as in Panel A were normalized to the value of each sample at Time = 0. There were statistically significant differences between the co-incubation and Jurkat cell samples, as indicated.

**Supplemental Figure 2:**
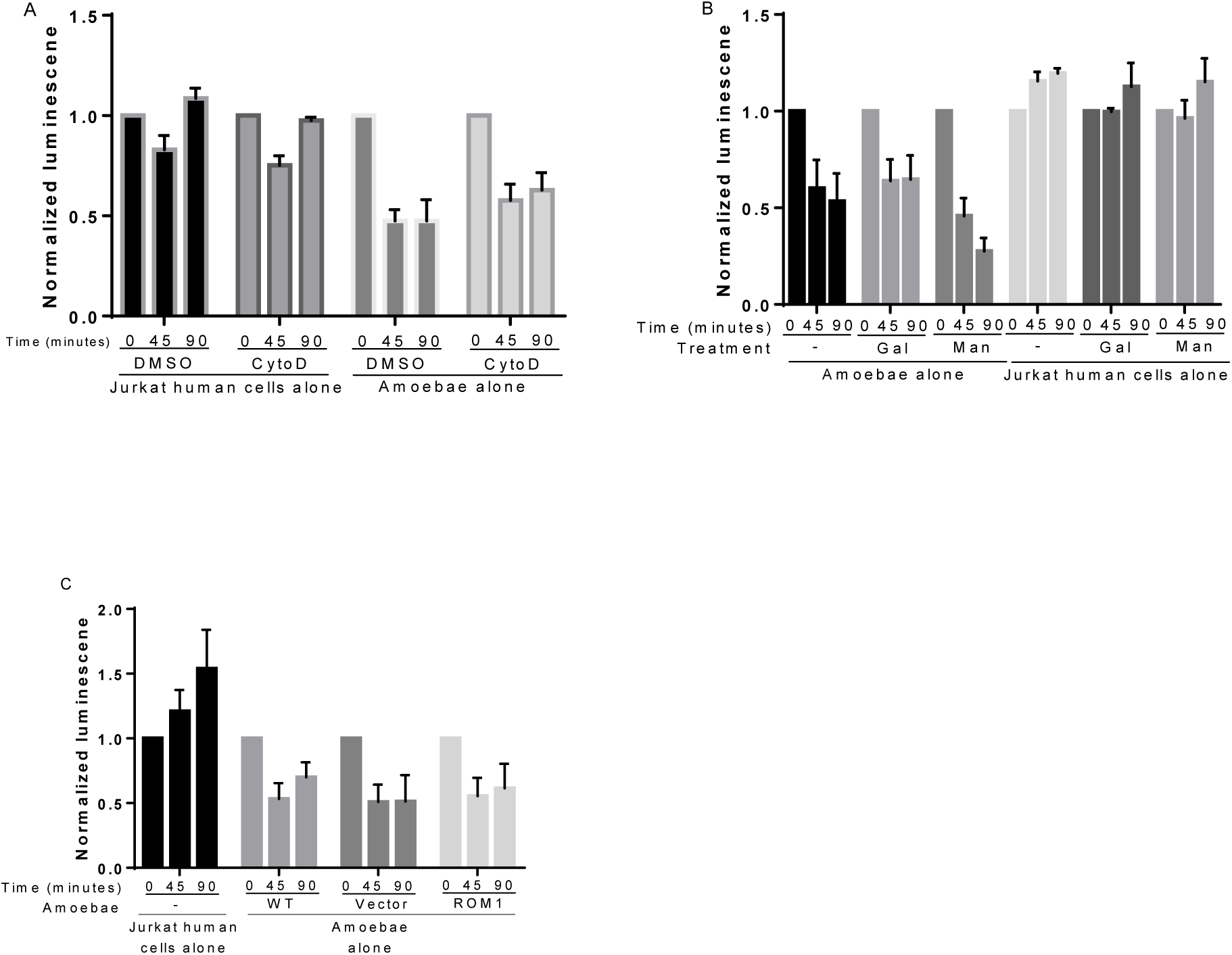
Additional controls for the CellTiterGlo assay data shown in Figure 2. **(A)** Amoebae and human Jurkat T cells were treated with Cytochalasin D or DMSO and viability was assayed using CellTiterGlo. Data from 4 independent experiments were normalized to the value of each sample at Time = 0. **(B)** Amoebae and Jurkat cells were incubated in media containing galactose, mannose, or no added sugar, and viability was assayed using CellTiterGlo. Data from 3 independent experiments were normalized to the value of each sample at Time = 0. **(C)** Amoebae were transfected with an EhRom1 knockdown plasmid, or a vector control plasmid. The viability of transfectants, wild-type non-transfected amoebae, and Jurkat cells was assayed using CellTiterGlo. Data from 3 independent experiments were normalized to the value of each sample at Time = 0.

**Supplemental Video 1: A dual-stain microscopy assay directly detects Jurkat cell killing by amoebae.** Amoebae and Hoechst-labeled human Jurkat T cells were co-incubated in the presence of SYTOX green. A representative video is shown, covering 20 minutes, captured at 2 frames/minute. Living human cells are labeled by Hoechst (blue), while dead human cells are labeled by both Hoechst and SYTOX green (green) and appear as turquoise in the merged video. Data are representative of 2 independent experiments.

**Supplemental Videos 2 and 3: A dual-stain microscopy assay directly detects Caco-2 cell killing by amoebae.** Amoebae and Hoechst-labeled human Caco-2 intestinal epithelial cells were co-incubated in the presence of SYTOX green. Representative videos are shown. Videos each follow the same field of cells and cover 20 minutes, captured at 1 frame/minute. Living human cells are labeled by Hoechst (blue), while dead human cells are labeled by both Hoechst and SYTOX green (green) and appear as turquoise in the merged video. Data are representative of 2 independent experiments.

## References

Baxt, L. A., Baker, R. P., Singh, U. and Urban, S. (2008). “An Entamoeba histolytica rhomboid protease with atypical specificity cleaves a surface lectin involved in phagocytosis and immune evasion.” Genes Dev 22(12): 1636–1646.

Baxt, L. A., Rastew, E., Bracha, R., Mirelman, D. and Singh, U. (2010). “Downregulation of an Entamoeba histolytica rhomboid protease reveals roles in regulating parasite adhesion and phagocytosis.” Eukaryot Cell 9(8): 1283–1293.

Bonora, M., Patergnani, S., Rimessi, A., De Marchi, E., Suski, J. M., Bononi, A., Giorgi, C., Marchi, S., Missiroli, S., Poletti, F., Wieckowski, M. R. and Pinton, P. (2012). “ATP synthesis and storage.” Purinergic Signal 8(3): 343–357.

Bracha, R. and Mirelman, D. (1984). “Virulence of Entamoeba histolytica trophozoites.” J. Exp. Med 160: 353–368.

Bracha, R., Nuchamowitz, Y., Leippe, M. and Mirelman, D. (1999). “Antisense inhibition of amoebapore expression in Entamoeba histolytica causes a decrease in amoebic virulence.” Molecular Microbiology 34(3): 463–472.

Debnath, A., Parsonage, D., Andrade, R. M., He, C., Cobo, E. R., Hirata, K., Chen, S., García-Rivera, G., Orozco, E., Martínez, M. B., Gunatilleke, S. S., Barrios, A. M., Arkin, M. R., Poole, L. B., McKerrow, J. H. and Reed, S. L. (2012). “A high-throughput drug screen for Entamoeba histolytica identifies a new lead and target.” Nature Med 18(6): 956–960.

Galluzzi, L., Bravo-San Pedro, J. M., Vitale, I., Aaronson, S. A., Abrams, J. M., Adam, D., Alnemri, E. S., Altucci, L., Andrews, D., Annicchiarico-Petruzzelli, M., Baehrecke, E. H., Bazan, N. G., Bertrand, M. J., Bianchi, K., Blagosklonny, M. V., Blomgren, K., Borner, C., Bredesen, D. E., Brenner, C., Campanella, M., Candi, E., Cecconi, F., Chan, F. K., Chandel, N. S., Cheng, E. H., Chipuk, J. E., Cidlowski, J. A., Ciechanover, A., Dawson, T. M., Dawson, V. L., De Laurenzi, V., De Maria, R., Debatin, K. M., Di Daniele, N., Dixit, V. M., Dynlacht, B. D., El-Deiry, W. S., Fimia, G. M., Flavell, R. A., Fulda, S., Garrido, C., Gougeon, M. L., Green, D. R., Gronemeyer, H., Hajnoczky, G., Hardwick, J. M., Hengartner, M. O., Ichijo, H., Joseph, B., Jost, P. J., Kaufmann, T., Kepp, O., Klionsky, D. J., Knight, R. A., Kumar, S., Lemasters, J. J., Levine, B., Linkermann, A., Lipton, S. A., Lockshin, R. A., Lopez-Otin, C., Lugli, E., Madeo, F., Malorni, W., Marine, J. C., Martin, S. J., Martinou, J. C., Medema, J. P., Meier, P., Melino, S., Mizushima, N., Moll, U., Munoz-Pinedo, C., Nunez, G., Oberst, A., Panaretakis, T., Penninger, J. M., Peter, M. E., Piacentini, M., Pinton, P., Prehn, J. H., Puthalakath, H., Rabinovich, G. A., Ravichandran, K. S., Rizzuto, R., Rodrigues, C. M., Rubinsztein, D. C., Rudel, T., Shi, Y., Simon, H. U., Stockwell, B. R., Szabadkai, G., Tait, S. W., Tang, H. L., Tavernarakis, N., Tsujimoto, Y., Vanden Berghe, T., Vandenabeele, P., Villunger, A., Wagner, E. F., Walczak, H., White, E., Wood, W. G., Yuan, J., Zakeri, Z., Zhivotovsky, B., Melino, G. and Kroemer, G. (2015). “Essential versus accessory aspects of cell death: recommendations of the NCCD 2015.” Cell Death Differ 22(1): 58–73.

Galluzzi, L., Vitale, I., Aaronson, S., Abrams, J., Adam, D., Agostinis, P., Alnemri, E., Altucci, L., Amelio, I., Andrews, D., Annicchiarico-Petruzzelli, M., Antonov, A., Arama, E., Baehrecke, E., Barlev, N., Bazan, N., Bernassola, F., Bertrand, M., Bianchi, K., Blagosklonny, MV, Blomgren, K., Borner, C., Boya, P., Brenner, C., Campanella, M., Candi, E., Carmona-Gutierrez, D., Ceccon, i. F., Chan, F., Chandel, N., Cheng, E., Chipuk, J., Cidlowski, J., Ciechanover, A., Cohen, G., Conrad, M., Cubillos-Ruiz, J., Czabotar, PE, D’Angiolella, V., Dawson, T., Dawson, V., De Laurenzi, V., De Maria, R., Debatin, K., DeBerardinis, RJ, Deshmukh, M, Di Daniele, N., Di Virgilio, F., Dixit, V., Dixon, S., Duckett, C., Dynlacht, B., El-Deiry, W., Elrod, J., Fimia, G., Fulda, S., García-Sáez, A., Garg, A., Garrido, C., Gavathiotis, E., Golstein, P., Gottlieb, E., Green, D., Greene, L., Gronemeyer, H., Gross, A., Hajnoczky, G., Hardwick, J., Harris, I., Hengartner, M., Hetz, C., Ichijo, H., Jäättelä, M., Joseph, B., Jost, P., Juin, P., Kaiser, W., Karin, M., Kaufmann, T., Kepp, O., Kimchi, A., Kitsis, R., Klionsky, D., Knight, R., Kumar, S., Lee, S., Lemasters, J., Levine, B., Linkermann, A., Lipton, S., Lockshin, R., López-Otín, C., Lowe, S., Luedde, T., Lugli, E., MacFarlane, M., Madeo, F., Malewicz, M., Malorni, W., Manic, G., Marine, J., Martin, S., Martinou, J., Medema, J., Mehlen, P., Meier, P., Melino, S., Miao, E., Molkentin, J., Moll, U., Muñoz-Pinedo, C., Nagata, S., Nuñez, G., Oberst, A., Oren, M., Overholtzer, M., Pagano, M., Panaretakis, T., Pasparakis, M., Penninger, J., Pereira, D., Pervaiz, S., Peter, M., Piacentini, M., Pinton, P., Prehn, J., Puthalakath, H., Rabinovich, G., Rehm, M., Rizzuto, R., Rodrigues, C., Rubinsztein, D., Rudel, T., Ryan, K., Sayan, E., Scorrano, L., Shao, F., Shi, Y., Silke, J., Simon, H., Sistigu, A., Stockwell, B., Strasser, A., Szabadkai, G., Tait, S., Tang, D., Tavernarakis, N., Thorburn, A., Tsujimoto, Y., Turk, B., Vanden Berghe, T., Vandenabeele, P., Vander Heiden, M., Villunger, A., Virgin, H., Vousden, K., Vucic, D., Wagner, E., Walczak, H., Wallach, D., Wang, Y., Wells, J., Wood, W., Yuan, J., Zakeri, Z., Zhivotovsky, B., Zitvogel, L., Melino, G. and Kroemer, G. (2018). “Molecular mechanisms of cell death: recommendations of the Nomenclature Committee on Cell Death 2018.” Cell Death Differ 25(3): 486–541.

Gilmartin, A., Ralston, K. and Petri, W. J. (2017). “Inhibition of Amebic Lysosomal Acidification Blocks Amebic Trogocytosis and Cell Killing.” mBio 8(4).

Huston, C. D., Houpt, E. R., Mann, B. J., Hahn, C. S. and Petri, W. A., Jr. (2001). “Caspase 3-dependent killing of host cells by the parasite Entamoeba histolytica.” Cellular Microbiology 2(6).

Kroemer, G., Galluzzi, L., Vandenabeele, P., Abrams, J., Alnemri, E. S., Baehrecke, E. H., Blagosklonny, M. V., El-Deiry, W. S., Golstein, P., Green, D. R., Hengartner, M., Knight, R. A., Kumar, S., Lipton, S. A., Malorni, W., Nuñez, G., Peter, M. E., Tschopp, J., Yuan, J., Piacentini, M., Zhivotovsky, B. and Melino, G. (2008). “Classification of cell death: recommendations of the Nomenclature Committee on Cell Death 2009.” Cell Death & Differentiation 16(1): 3–11.

Leist, M., Single, B., Castoldi, A. F., Kuhnle, S. and Nicotera, P. (1997). “Intracellular Adenosine Triphosphate (ATP) Concentration: A Switch in the Decision Between Apoptosis and Necrosis.” Journal of Experimental Medicine 185(8).

Li, E., Stenson, W. F., Kunz-Jenkins, C., Swanson, P. E., Duncan, R. and Stanley, S. L. J. (1994). “Entamoeba histolytica Interactions with Polarized Human Intestinal Caco-2 Epithelial Cells.” Infection and Immunity 62(11): 5112–5119.

Lidell, M. E., Moncada, D. M., Chadee, K. and Hansson, G. C. (2006). “Entamoeba histolytica cysteine proteases cleave the MUC2 mucin in its C-terminal domain and dissolve the protective colonic mucus gel.” Proc Natl Acad Sci U S A 103(24): 9298–9303.

Marie, C. S., Verkerke, H. P., Paul, S. N., Mackey, A. J., Petri, W. A. Jr. (2012). “ Leptin protects host cells from Entamoeba histolytica cytotoxicity by a STAT3-dependent mechanism.” Infect Immun 80(5): 1934–1943.

Miller, H. W., Suleiman, R. L. and Ralston, K. S. (2019). “Trogocytosis by Entamoeba histolytica Mediates Acquisition and Display of Human Cell Membrane Proteins and Evasion of Lysis by Human Serum.” mBio 10(2).

Morf, L., Pearson, R. J., Wang, A. S. and Singh, U. (2013). “Robust gene silencing mediated by antisense small RNAs in the pathogenic protist Entamoeba histolytica.” Nucleic Acids Research 41(20): 9424–9437.

Petri, W. A., Jr., Haque, R. and Mann, B. J. (2002). “The bittersweet interface of parasite and host: lectin-carbohydrate interactions during human invasion by the parasite Entamoeba histolytica.” Annu Rev Microbiol 56: 39–64.

Ralston, K. S. (2015). “Taking a bite: Amoebic trogocytosis in Entamoeba histolytica and beyond.” Curr Opin Microbiol 28: 26–35.

Ralston, K. S. and Petri, W. A., Jr (2011). “Tissue destruction and invasion by Entamoeba histolytica.” Trends Parasitol 27(6): 254–263.

Ralston, K. S., Solga, M. D., Mackey-Lawrence, N. M., Somlata, Bhattacharya, A. and Petri, W. A., Jr. (2014). “Trogocytosis by Entamoeba histolytica contributes to cell killing and tissue invasion.” Nature 508(7497): 526–530.

Ravdin, J. I., Croft, B. Y. and Guerrant, R. L. (1980). “Cytopathogenic mechanisms of Entamoeba histolytica.” J Exp Med 152(2): 377–390. (a)

Ravdin, J. I. and Guerrant, R. L. (1980). “Studies on the cytopathogenicity of Entamoeba histolytica.” Arch. Invest. Med. (Mex) 11: 123–128. (b)

Ravdin, J. I. and Guerrant, R. L. (1981). “Role of adherence in cytopathogenic mechanisms of Entamoeba histolytica. Study with mammalian tissue culture cells and human erythrocytes.” J Clin Invest 68(5): 1305–1313.

Ravdin, J. I., Murphy, C. F., Guerrant, R. L. and Long-Krug, S. A. (1985). “Effect of Antagonists of Calcium and Phospholipase A on the Cytopathogenicity of Entamoeba histolytica.” The Journal of Infectious Diseases 152(3): 542–549.

Reed, S., Ember, J., Herdman, D., DiScipio, R., Hugli, T. and Gigli, I. (1995). “The extracellular neutral cysteine proteinase of Entamoeba histolytica degrades anaphylatoxins C3a and C5a.” J Immunol 155(1): 266–274.

Riss, T. and Niles, A., Moravec, R, Karassina, N, Vidugiriene, J (2019). Cytotoxicity Assays: In Vitro Methods to Measure Dead Cells. Bethesda (MD), Eli Lilly & Company and the National Center for Advancing Translational Sciences.

Saffer, L. D. and Petri, W. A. (1991). “Role of the galactose lectin of Entamoeba histolytica in adherence-dependent killing of mammalian cells.” Infection and Immunity 59(12): 4681–4683.

Seydel, K. and Stanley, S. J. (1998). “Entamoeba histolytica induces host cell death in amebic liver abscess by a non-Fas-dependent, non-tumor necrosis factor alpha-dependent pathway of apoptosis.” Infect Immun. 66(6): 2980–2983.

Tang, H., Tang, H., Mak, K., Hu, S., Wang, S., Wong, K., Wong, C., Wu, H., Law, H., Liu, K., Talbot, C. J., Lau, W., Montell, D. and Fung, M. (2012). “Cell survival, DNA damage, and oncogenic transformation after a transient and reversible apoptotic response.” Mol Biol Cell 23(12): 2240–2252.

Teixeira, J. E., Sateriale, A., Bessoff, K. E. and Huston, C. D. (2012). “Control of Entamoeba histolytica Adherence Involves Metallosurface Protease 1, an M8 Family Surface Metalloprotease with Homology to Leishmanolysin.” Infect Immun 80(6): 2165–2176.

Thibeaux, R., Dufour, A., Roux, P., Bernier, M., Baglin, A. C., Frileux, P., Olivo-Marin, J. C., Guillen, N. and Labruyere, E. (2012). “Newly visualized fibrillar collagen scaffolds dictate Entamoeba histolytica invasion route in the human colon.” Cell Microbiol 14(5): 609–621.

Tillack, M., Nowak, N., Lotter, H., Bracha, R., Mirelman, D., Tannich, E. and Bruchhaus, I. (2006). “Increased expression of the major cysteine proteinases by stable episomal transfection underlines the important role of EhCP5 for the pathogenicity of Entamoeba histolytica.” Molecular and Biochemical Parasitology 149(1): 58–64.

